# Design, Development and Validation of New Fluorescent Strains for Studying Oral Streptococci

**DOI:** 10.1101/2025.01.14.632972

**Authors:** Daniel I. Peters, Iris J. Shin, Alyssa N. Deever, Justin R. Kaspar

**Affiliations:** Division of Biosciences, The Ohio State University College of Dentistry, Columbus, Ohio

**Author notes:** Corresponding author **Mailing address:** Division of Biosciences, The Ohio State University, College of Dentistry, 305 W. 12^th^ Avenue, Postle Hall Rm 4185, Columbus, OH 43210.

## Abstract

Bacterial strains that are genetically engineered to constitutively produce fluorescent proteins have aided our study of bacterial physiology, biofilm formation, and interspecies interactions. Here, we report on the construction and utilization of new strains that produce the blue fluorescent protein mTagBFP2, the green fluorescent protein sfGFP, and the red fluorescent protein mScarlet-I3 in species *Streptococcus gordonii, Streptococcus mutans*, and *Streptococcus sanguinis*. Gene fragments, developed to contain the constitutive promoter P*veg*, the fluorescent gene of interest as well as *aad9*, providing resistance to the antibiotic spectinomycin, were inserted into selected open reading frames on the chromosome that were both transcriptionally silent and whose loss caused no measurable changes in fitness. All strains, except for sfGFP in *S. sanguinis*, were validated to produce a detectable and specific fluorescent signal. Individual stains, along with extracellular polymeric substances (EPS) within biofilms, were visualized and quantified through either widefield or super-resolution confocal microscopy approaches. Finally, to validate the ability to perform single cell-level analysis using the strains, we imaged and analyzed a triculture mixed-species biofilm of *S. gordonii, S. mutans*, and *S. sanguinis* grown with and without addition of human saliva. Quantification of the loss in membrane integrity using a SYTOX dye revealed that all strains had increased loss of membrane integrity with water or human saliva added to the growth media, but the proportion of the population stained by the SYTOX dye varied by species. In all, these fluorescent strains will be a valuable resource for the continued study of oral microbial ecology.

## INTRODUCTION

Bacteria of the genus *Streptococcus* colonize almost every location in the human body, are able to persist in a commensal state, or at times, are a cause of some of our species’ most common diseases, including pneumonia, pharyngitis, meningitis, urinary tract infections, and tooth decay. Streptococci are the primary colonizers at different sites in the oral cavity (1–3), including the soft tissues of the tongue and gingiva, as well as the hard tissues of tooth enamel (4–8). Oral streptococci adhere to theses surfaces, and other oral bacteria, in part through direct binding mediated by the actions of encoded adhesins, as well as indirectly through the development of biofilms, consisting both of bacterial cells as well as extracellular polymeric substances (EPS) including extracellular DNA (eDNA) and polysaccharides (9, 10). In recent years, there has been a renewed interest in studying the physiology of streptococci either in association with the host or in microbial mixed-species communities (11, 12). In addition, the coupling of advanced imaging with other molecular microbiology techniques have continued the need of bacterial species and/or strain labeling/tagging applications. There is a constant need to both evaluate and optimize newly developed methods, as well as the incorporation of new probe variations, stains, and fluorescent proteins.

Several strains of oral streptococci have been cloned to contain fluorescent genes from which expression is driven by a constitutive promoter, intended for use in microscopy as well as other applications (13–15). Often these strains may not be suitable for their desired use, either due to their lack of a strong and/or specific fluorescent signal, and/or due to the instability of their vector within the host strain that contains the fluorescent constructs. In addition, the current suite of available fluorescent strains is limited in variety to either green fluorescent protein (GFP) or red fluorescent protein (RFP), which may prevent the use of other fluorescent cells and/or stains with similar excitation/emission (Ex/Em) spectral overlap.

In the last several years, there has been continued development of new fluorescent protein variants that improve on characteristics such as brightness, stability, and maturation time (16). In addition, the availability and affordability to design and synthesize gene fragments using synthetic biology approaches have lowered the barriers towards developing, cloning, and assaying new bacterial strain variants that produce a wider range of fluorescent proteins (17). Here in this study, we took advantage of these progresses to design, develop and experimentally validate strains of *Streptococcus gordonii, Streptococcus mutans* and *Streptococcus sanguinis* that produce mTagBFP2, sfGFP and mScarlet-I3 – blue, green and red fluorescent proteins. We show how these strains can be utilized to further explore mixed-species interactions, biofilm formation and development, as well as combined with other labeling approaches to quantify species’ behaviors on a single-cell level using super-resolution confocal microscopy. We believe that both these resources themselves, as well as the general approach that can be mirrored in other *Streptococcus* and bacterial species, will be valuable contributions to the fields of oral microbiology and biofilm research and will add to our toolbox of resources that will allow further exploration in how bacterial communities form, and the interactions between microbes contained within them.

## RESULTS

### Chromosomal insertion into sites that lack transcriptional activity lead to no change in bacterial fitness

A previous iteration of promoter-fluorescent gene constructs were cloned into the *E. coli*-streptococcal shuttle vector pDL278, which allowed transformation of the same plasmid into multiple different species of oral streptococci (13). While this was advantageous in utilizing the same construct across multiple strains and thereby reducing cloning efforts, this approach also carried disadvantages, such as usage of antibiotics during strain growth to provide selective pressure against plasmid segregational instability and higher fluorescent gene expression noise due to variability in plasmid copy number between individual cells that was observed by the wide range of fluorescent intensities across a population. As our end goal was to utilize fluorescent strains for single cell applications, we took these considerations into account and chose to design synthetic gene fragments with the end purpose of integrating them into the host strain’s bacterial chromosome.

To determine the optimal site(s) for inserting a promoter-fluorescent gene construct without distribution to the normal cell transcriptome and overall physiology, we first reviewed previously acquired RNA-Seq datasets where *S. mutans, S. gordonii* and *S. sanguinis* had been grown in TYG medium to mid exponential growth phase (18). We sorted RNA read counts from low to high across all called genome features within files produced from htseq-count after read mapping to the genome, and selected a site in each species that had zero to very low (∼10) read counts in a hypothetical gene with no known determined function **(Figure 1A)**. This included SMU.1155 in *S. mutans* UA159, a hypothetical gene that overlaps with SMU.1154c and SMU.1156c and has no homology to genes in other oral streptococci species, SGO_2075 in *S. gordonii* DL1 that is described as a plasmid replication protein and appears to be in a region of the chromosome that are gene remnants of a prophage (e.g., SMU_2076 is a phage integrase) and an area that has previously been used as a chromosomal integration site (19), and SSA_20230 in *S. sanguinis* SK36, a hypothetical gene with 450+ bp of intergenic region surrounding it in both the 5’ and 3’ direction.

**Figure 1.**
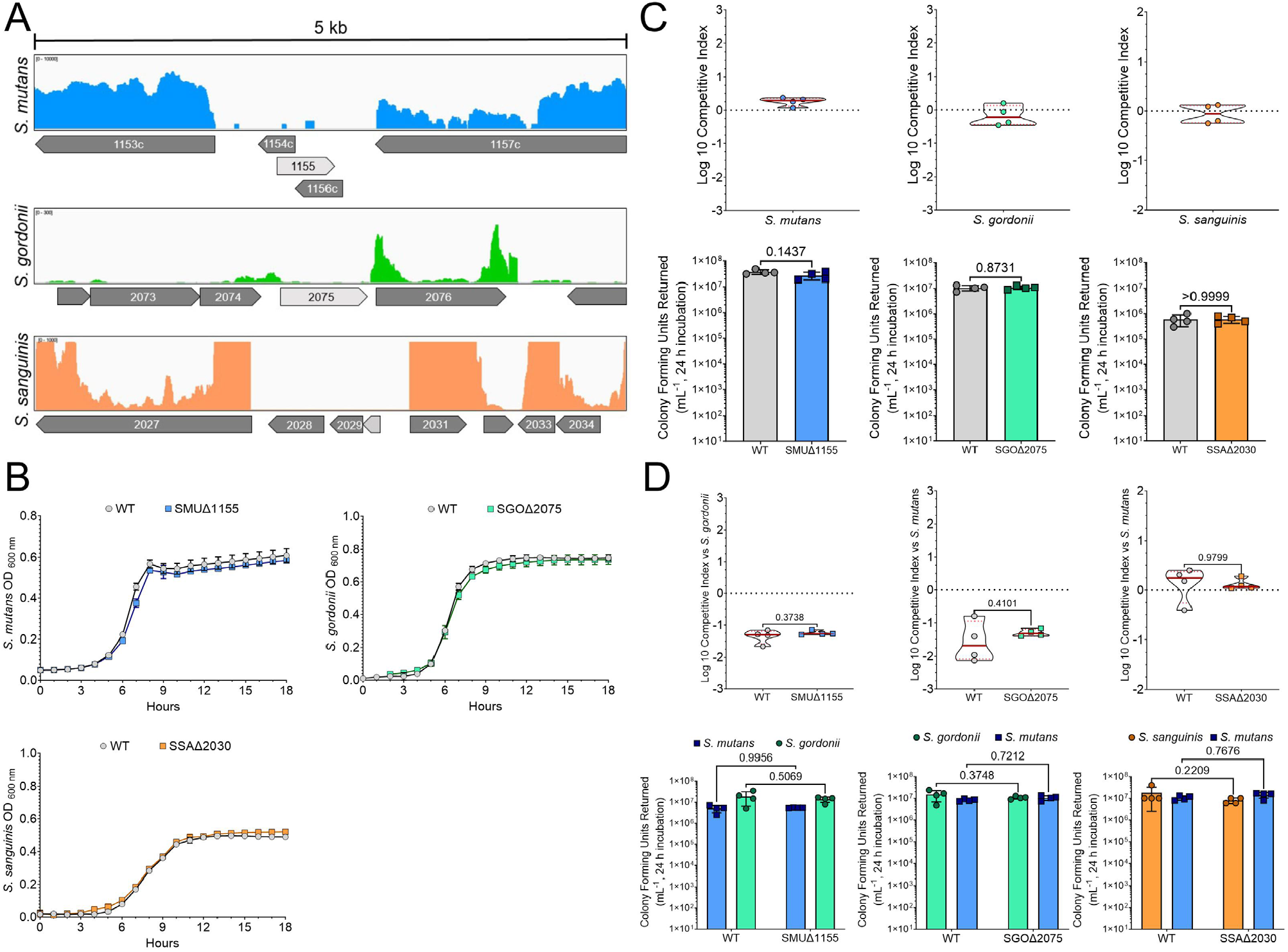
Location of chromosomal fluorescent gene insertion does not affect strain fitness. **(A)** Distribution of RNA-Seq reads surrounding chosen chromosomal site to insert fluorescent gene fragment in either *S. mutans* (blue top panel), *S. gordonii* (green middle panel), or *S. sanguinis* (bottom orange panel). Each colored line/peak represents a single RNA read, and thus active transcription, at the time of cell harvest. The gene that corresponds to the insertion site is shown in light gray, while surrounding genes are shown in dark gray and indicated by their gene number, if space allows. Each selected gene (SMU_1155, SGO_2075 and SSA_2030) shows little to no active transcription (in this culturing condition, TYG). The scale of reads within each region is noted in the top left of each panel. **(B)** Growth curve comparison of the wild-type (WT, gray circles) strain and insertion site mutant (colored squares) in TYG medium. Cultures were grown for 18 h with an optical density reading every hour. **(C)** Log 10 competitive index (top graphs) between the wild-type and insert site mutant. A competitive index near or at 0 indicates no fitness advantage present for either strain. Final CFU counts from the competitive index experiment between the WT and insert site mutant (bottom graphs). Statistical analysis is shown using an unpaired t-test with Welch’s correction. n = 4. **(D)** Log 10 competitive index (top graphs) between the WT or insert site mutant with another oral *Streptococcus* species (*S. gordonii* for *S. mutans, S. mutans* for *S. gordonii* and *S. sanguinis*). There is no significant fitness changes between the WT and insert site mutant within these mixed-species cocultures. Final CFU counts from the competitive index experiment (bottom graphs). Statistical analysis is shown using a two-way analysis of variance (ANOVA) with Sidak’s multiple comparisons test. n = 4.

To ensure that insertion into these areas did not impact strain fitness, we first constructed mutants via allelic replacement of the selected genome feature with *ermB*, conferring erythromycin resistance, using a PCR ligation mutagenesis approach (20). Growth of the resulting mutant strains was comparable to the wild-type (WT; parental) strain in TYG medium **(Figure 1B)**. In addition, we also performed competitive index experiments of the mutants in coculture with either the WT **(Figure 1C)**, or an oral streptococci competitor (*S. gordonii* for *S. mutans*, and *S. mutans* for *S. gordonii* and *S. sanguinis*) as the intended use of these strains is for mixed-species biofilm imaging **(Figure 1D)**. In coculture against the WT, the Log 10 competitive index was at or near zero, suggesting no competitive advantage for either strain over a 24 h period. This was also confirmed by comparing the colony forming units (CFUs) returned after 24 h from the competitive index experiments that displayed no significant differences. In coculture against an oral streptococci competitor, both the WT and insertion site mutants returned similar competitive indexes and final CFUs for both species present in the coculture. Therefore, deletion and/or insertion into these areas does not influence strain growth or fitness in cocultures, at least when grown in TYG medium.

### Strains expressing fluorescent genes show similar growth and biofilm formation profiles

We designed a synthetic promoter-fluorescent gene construct to be cloned into our selected site(s) **(Figure 2A, Supplemental Figures 1-3)**. We chose fluorescent genes *mtagbfp2*, coding for a blue fluorescent protein with excitation/emission (Ex/Em) maxima at 399/454 (21), *sfgfp* which we have previously used in oral streptococci imaging applications (13), and *mscarlet-I3*, a recently developed red fluorescent protein that has undergone several rounds of targeted and random mutagenesis to exhibit higher brightness and faster maturation than previous versions of mScarlet (22). The promoter P*veg*, with advantageous characteristics described elsewhere (13), drives expression of both the fluorescent gene and *aad9* that confers spectinomycin resistance to be used during transformation selection while undergoing cloning, as well as other assay(s) where selection may be desired (such as competitive index experiments described above). Finally, a strong transcriptional terminator L3S2P21 (23) was included so that read through from the P*veg* promoter into surrounding regions would be minimal. The construct is surrounded on both sides by BamHI restriction sites for PCR ligation mutagenesis cloning applications.

**Figure 2.**
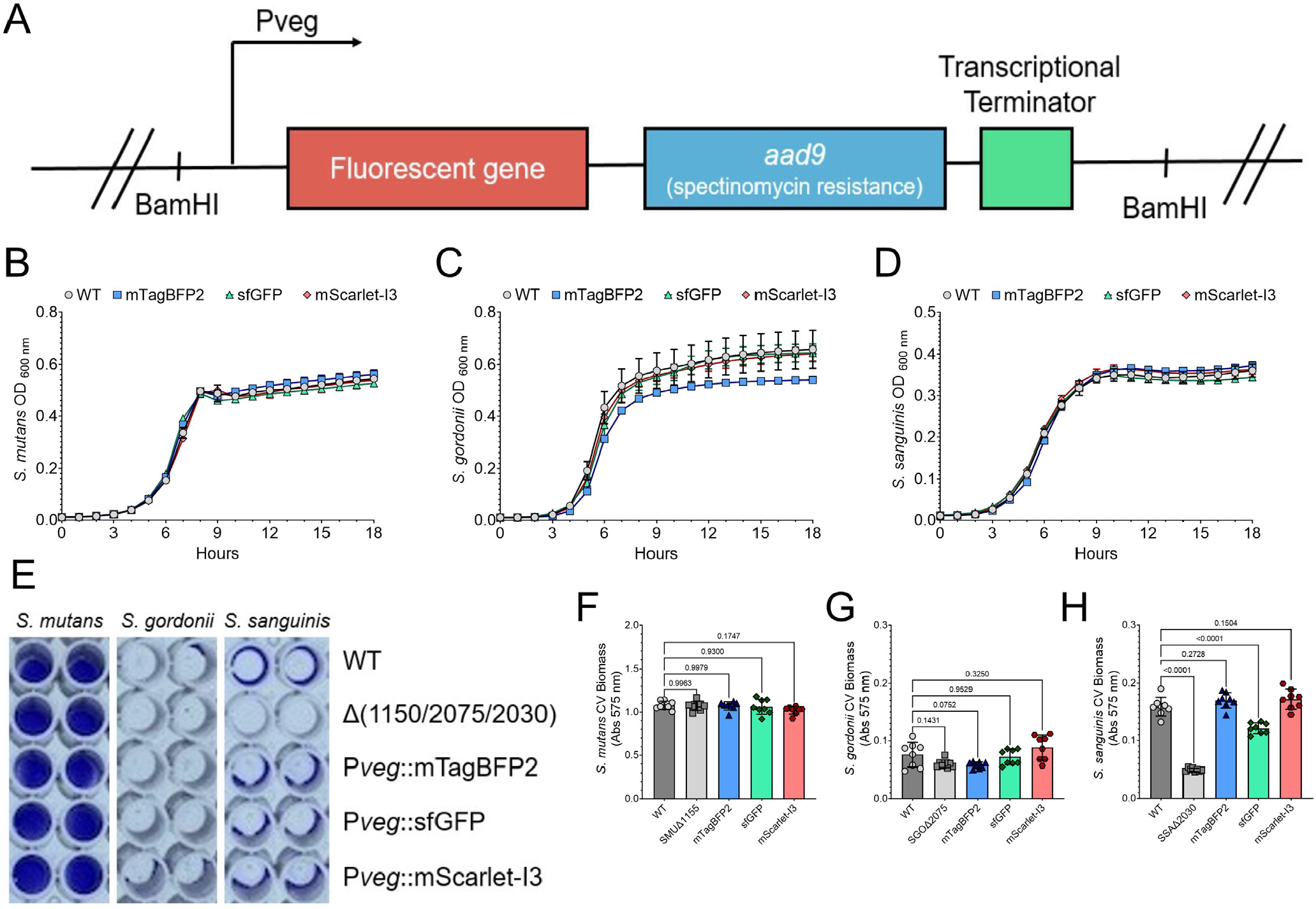
Growth and biofilm formation is comparable to wild-type for the cloned fluorescent strains. **(A)** Diagram of the fluorescent gene fragment that was inserted into the chromosome at the insertion site. Both of the 5’ and 3’ ends contain a BamHI restriction enzyme site for PCR ligation mutagenesis. The promoter, P*veg*, drives expression of the fluorescent gene (*mtagbfp2, sfgfp* or *mscarlet-I3*), and *aad9* provides resistance for spectinomycin, used for cloning and other assay selection. A strong transcriptional terminator (L3S2P21) is included after both coding genes. Growth curve comparison of the wild-type (WT, gray circles) strain and fluorescent strains mTagBFP2 (blue squares), sfGFP (green triangles) and mScarlet-I3 (red diamonds) in either **(B)** *S. mutans*, **(C)** *S. gordonii* or **(D)** *S. sanguinis*. Cultures were grown in TYG medium for 18 h with an optical density reading every hour. **(E)** Representative image of a crystal violet (CV) biofilm biomass assay where strains (species listed on top and strains down right y-axis) were grown in TYG supplemented with 5 mM sucrose for 24 h. Quantification of the CV assay shown in E for **(F)** *S. mutans*, **(G)** *S. gordonii* or **(H)** *S. sanguinis* strains. Statistical analysis is shown using an ordinary one-way analysis of variance (ANOVA) with Dunnett’s multiple comparisons test. n = 8.

Synthetic gene fragment constructs, one for each fluorescent gene, were cloned and transformed into *S. mutans, S. gordonii* and *S. sanguinis* at the previously described chromosomal insertion site. Resulting strains were first assayed for their growth properties in comparison to the WT strain **(Figure 2B-2D)**, as well as biofilm biomass accumulation using the crystal violet (CV) biofilm assay **(Figure 2E-2H)**. In terms of growth, only Pveg::mTagBFP2 in *S. gordonii* displayed a growth profile that differed from the WT, and this was mainly as the strain reached late exponential to stationary growth phase. All other strains were similar to their parental control. For biofilm biomass accumulation over a 24 h period in TYGS medium (i.e., TYG containing 5 mM sucrose), all strains of *S. mutans* and *S. gordonii* did not show any significant differences in biofilm formation. However, two strains of *S. sanguinis*, the *ermB* insertion site mutant (SSAΔ2030) and Pveg::sfGFP, showed significant reduction in biomass accumulation by CV. In general, almost all fluorescent strains constructed in this study displayed characteristic growth and biofilm formation properties that represent their parental strain for forthcoming single- and mixed-species biofilm assays.

### Characterization of fluorescent intensity and maturation time across three oral streptococci species

Our next objective was to determine (i) how well each strain produced a detectable fluorescent signal, (ii) whether that signal was specific for the emission range (i.e., channel) already described for that fluorescent protein, and (iii) the maturation time required for each protein to emit a detectable signal. To this end, we monitored the detectable fluorescent intensity of each strain over 18 h of growth in TYG using multiple Ex/Em settings that would correspond to detection in a DAPI, FITC, or Texas Red channel **(Figure 3A-3C)**. For example, the detected fluorescent intensity for all three *S. mutans* strains using Ex/Em settings that correspond to the DAPI channel showed a highly specific signal for our constructed mTagBFP2 strain that could be quantified over background after 9 h of growth, as expected. In fact, all three *S. mutans* strains showed highly specific signal for their respective channel (sfGFP for FITC, mScarlet-I3 for Texas Red). This was also the case for all three strains of *S. gordonii*, although the sfGFP strain did show early (between 3 – 7 h) fluorescent intensity in the DAPI channel. Both of the mTagBFP2 and mScarlet-I3 displayed detectable intensity over background around the 9 h mark of growth, while sfGFP was detectable earlier (∼5-6 h of growth). However, these trends did not track in *S. sanguinis*, where mTagBFP2 displayed higher intensity after 15 h of growth, sfGFP was not detectable over background, and while intensity of mScarlet-I3 was measurable over time, it did not drastically rise at the 9 h mark as seen in *S. mutans* and *S. gordonii*. Therefore, the maturation time required and production of measurable fluorescent intensity utilizing the same synthetic gene fragment is species-specific, a similar observation we previously made with fluorescent vectors (13).

**Figure 3.**
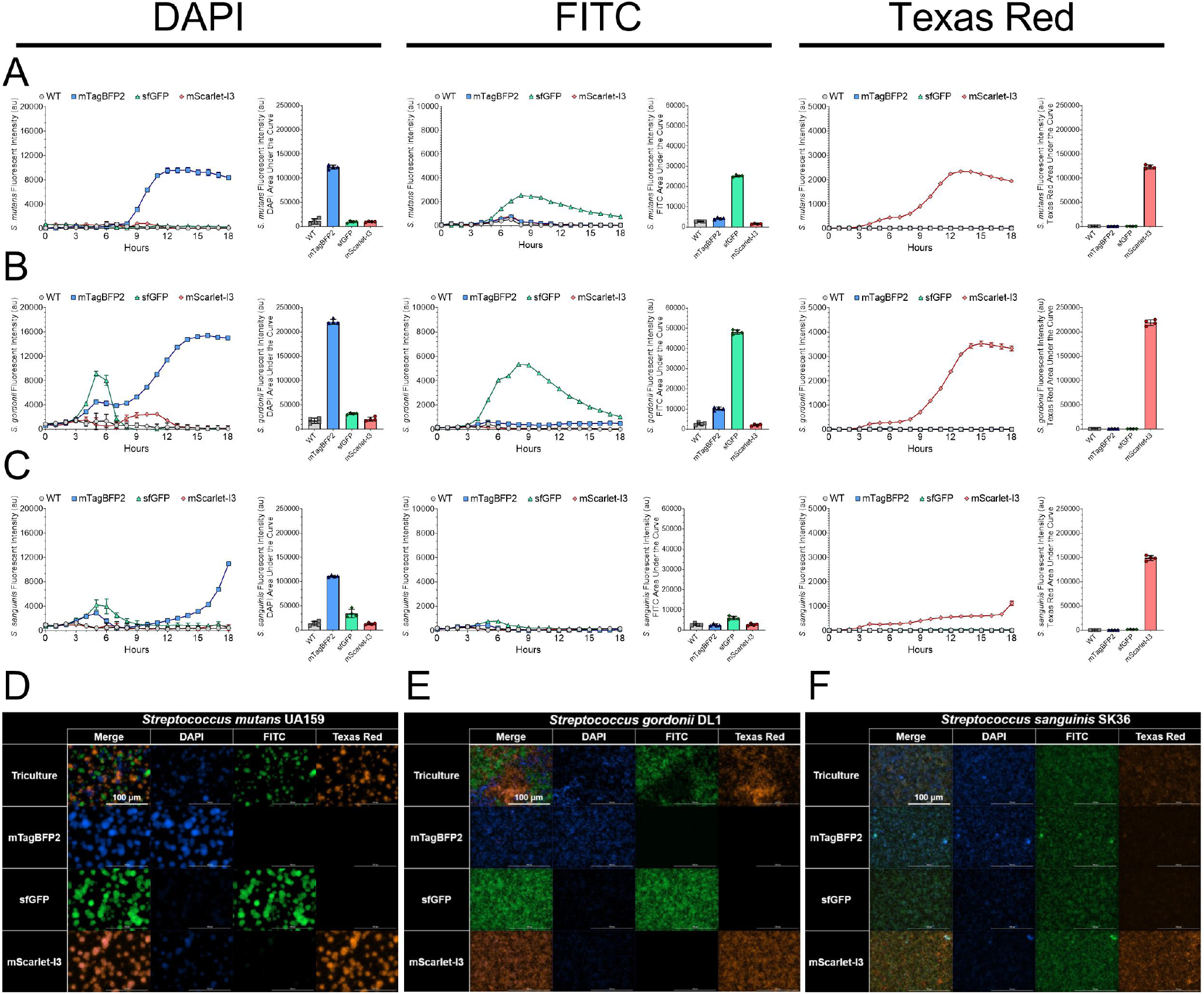
Fluorescent protein maturation and intensity production between strains is species specific. Relative fluorescent intensity (arbitrary units, a.u.) over 18 h for either the wild-type (WT, gray circles) strain or fluorescent strains mTagBFP2 (blue squares), sfGFP (green triangles) and mScarlet-I3 (red diamonds) in either **(A)** *S. mutans*, **(B)** *S. gordonii* or **(C)** *S. sanguinis*. The left column represents intensities in the DAPI channel (Ex 399 / Em 455), the middle column represents the FITC channel (Ex 485 / Em 528), and the right column the Texas Red channel (Ex 550 / Em 590). The left graph displays fluorescent intensity over time, highlighting different fluorescent protein maturation times in each strain, while the right graph shows the cumulative area under the curve (AUC) for the graph on the left (i.e., cumulative fluorescent intensity over 18 h). Maximum intensity, 40x Z-projections of triculture and individual monoculture biofilms for the three fluorescent **(D)** *S. mutans*, **(E)** *S. gordonii*, or **(F)** *S. sanguinis* strains with image captures within each specific channel (DAPI, FITC and Texas Red) as well as a merged image. Biofilms were grown for 24 h in TYG supplemented with 5 mM sucrose prior to imaging. Scale bar (100 µm) is shown in top left image.

We also visualized fluorescent production of each strain individually in monoculture across all channels, as well as in a triculture with all three strains together **(Figure 3D-3F)**. For both *S. mutans* and *S. gordonii*, each strain produced an easily detectable signal that was specific for its intended channel, similar to the plate reader experiments described above. It should be noted that mScarlet-I3 did produce a detectable DAPI signal, along with a weaker sfGFP signal – both of which would be manageable through adjustment of instrument settings such as a reduction in exposure time to acquire the image (settings were set higher than normal in this experiment than would be otherwise used so that full signal acquisition was achieved). However in *S. sanguinis*, both of the mTagBFP2 and mScarlet-I3 strains showed signal in the FITC channel, and intensity from sfGFP was noticeably weaker than in *S. mutans* or S. *gordonii*, again similar to data captured by the plate reader experiments. Even with these noted issues, the mTagBFP2 or mScarlet-I3 strains of *S. sanguinis* can still be utilized knowing that image acquisition settings may need to be specifically tuned to account for lack of signal intensity.

In addition to widefield microscopy, we also visualized either *S. mutans* or *S. gordonii* biofilm tricultures of all three strains by super-resolution confocal microscopy **(Figure 4)**. Here, all three strains again produced signals specific for their respective channel such that the spatial arrangement of all three strains could be clearly observed individually in the triculture. This is best seen in the *S. mutans* triculture, where strain-specific *S. mutans* microcolonies could be differentiated, suggesting that each microcolony originated from a single strain, and not formed from a combination of different *S. mutans* strains. We were also able to observe the location, arrangement, and quantity of extracellular glucan polysaccharides within the biofilms, with an extra channel (Alexa647; CY5) available for imaging. Detectable glucans were the most intense in the center of the *S. mutans* microcolony structures. In contrast, very little glucans were detectable in the *S. gordonii* triculture, as expected.

**Figure 4.**
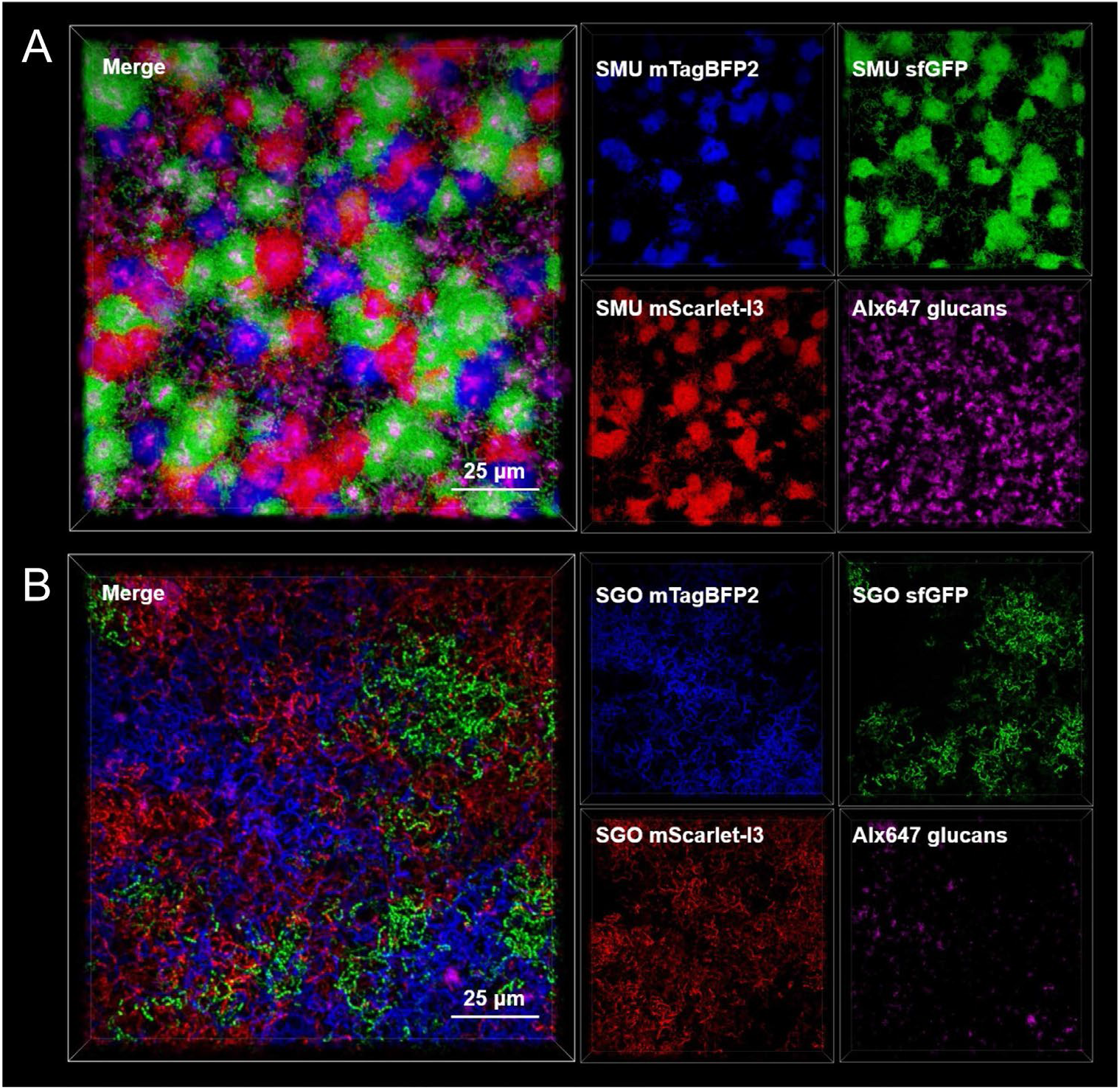
Super-resolution confocal imaging of single species tricultures. Maximum intensity, 100x 3D models of super-resolution confocal-captured biofilm images oriented from the top down (Z+) of either **(A)** *S. mutans* (SMU) or **(B)** *S. gordonii* (SGO) strain triculture. The merged image is shown on the left, with individual channels on the right (DAPI channel blue, FITC channel green, Texas Red channel red, and CY5 channel pink). Biofilms were grown for 24 h in TYG supplemented with 5 mM sucrose prior to imaging. Scale bar (25 µm) is shown in the bottom right of the merged image. Alexa Fluor 647-labeled dextran was added during bacterial strain inoculation and shows glucan production within either triculture. Images are 127 μm (L) x 127 μm (W) x 30 μm (H).

### Individual strains and biofilm matrix components can be visualized and quantified from mixed-species biofilms

Having verified that these strains produce a strain-specific fluorescent signal, we next imaged a coculture of *S. mutans* sfGFP and *S. gordonii* mTagBFP2 **(Figure 5A)**. In addition, we visualized biofilm extracellular matrix components through labeling eDNA using a dsDNA-specific antibody (24) as well as the aforementioned glucans (25, 26). All four strains/components could be observed and quantified within their respective channel. *S. mutans* growth was restrictive to their microcolony structures, while *S. gordonii* formed a thin layer of cells in spaces not occupied by the *S. mutans* microcolonies. This is similar to previous observations we and others have made for imaged oral streptococci biofilms. When observing biofilm matrix components, a majority of the labeled eDNA and glucans were contained within, yet not limited to, *S. mutans* microcolony structures. Dual labeling of both eDNA and glucans could be observed both within *S. mutans* microcolonies, as well as associated with *S. gordonii*. While both bacterial strains made up 79% of the biomass volume (58±4% for S. *mutans* and 21±2% for *S. gordonii*), 21% was made up of extracellular matrix components, with glucans being 14±1% and eDNA 7±1% **(Figure 5B)**. Finally, as each biofilm component could be segregated into individual channels, we viewed each component with z-depth coding to give perspectives on the thickness of each component **(Figure 5C)**. This confirmed that *S. gordonii* forms a thinner layer of cells (5 – 10 μm) with *S. mutans* microcolonies towering above (10 – 20 μm). Yet, both eDNA and glucans can extend over the height of the bacterial biofilm, with glucans being the thickest measured component (20 – 25 μm). In all, this experiment validates usage of these strains in mixed-species settings to further interrogate the relationship between individual bacterial species and resulting biofilm matrix components during interspecies interactions.

**Figure 5.**
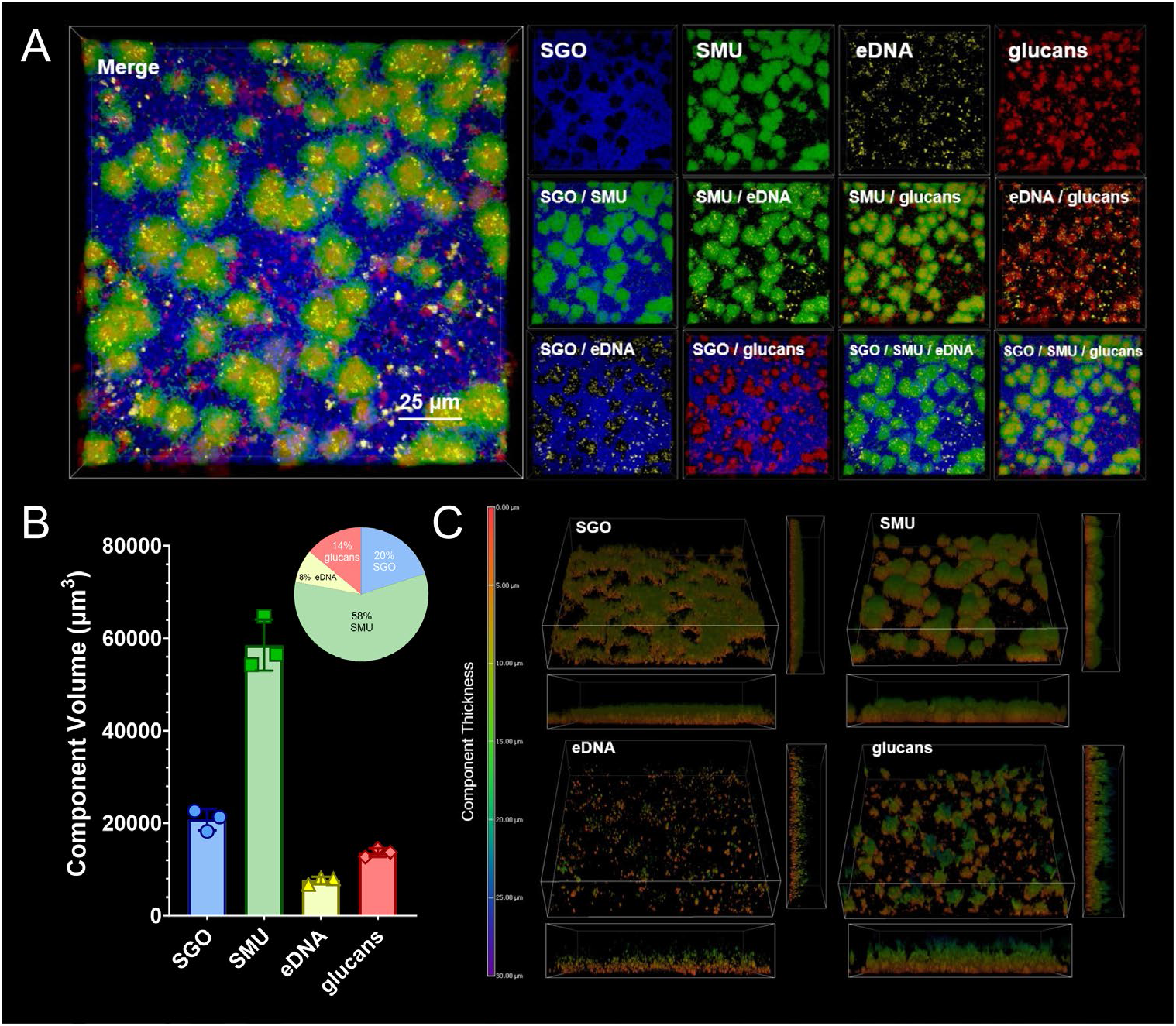
Super-resolution confocal imaging of S. mutans and S. gordonii coculture. **(A)** Maximum intensity, 100x 3D model of a super-resolution confocal-captured biofilm image oriented from the top down (Z+) of *S. gordonii* (SGO; mTagBFP2, blue) and *S. mutans* (SMU; sfGFP, green) coculture. Biofilm matrix components eDNA (yellow; α-dsDNA, 35I9 DNA) and glucans (red; Alexa Fluor 647-labeled dextran) were also imaged and shown. The merged image is shown on the left, with individual and combined channels on the right. Biofilms were grown for 24 h in TYG supplemented with 5 mM sucrose prior to imaging. Scale bar (25 µm) is shown in the bottom right of the merged image. Images are 127 μm (L) x 127 μm (W) x 30 μm (H). **(B)** Bar and pie graph of calculated component volumes of the biofilms shown in A. The bar graph displays the quantified volumes, while the pie graph shows the percentage of each component of the total biofilm volume. Three independent images/areas were quantified from the same biofilm sample. **(C)** Z-depth coded perspective 3D models of the biofilm image shown in A, broken out into individual components/channels. Depth coding scale corresponding to component thickness is shown on the left in microns (0-30 μm). Red-orange represents little to no thickness, greens represent medium thickness and blue-purple represent high thickness. X- and y-side perspectives are shown on the bottom and right side of the tilted Z+ perspective image.

### Mixed-species biofilms grown in human saliva display species-specific cell death

Previously our group has shown that oral streptococci modify their behaviors when grown with human saliva (27). To better understand how inclusion of saliva into the growth media alters biofilm architecture and species-level distribution, we inoculated triculture biofilms of *S. sanguinis* mTagBFP2, *S. mutans* sfGFP and *S. gordonii* mScarlet-I3 into TYG-, TYG-H_2_O or TYG-Saliva supplemented with 5 mM sucrose and imaged the biofilms with super-resolution confocal microscopy 24 h later **(Figure 6A)**. Similar to the coculture biofilm discussed above, all three strains could be observed and quantified individually within this experimental setup.

**Figure 6.**
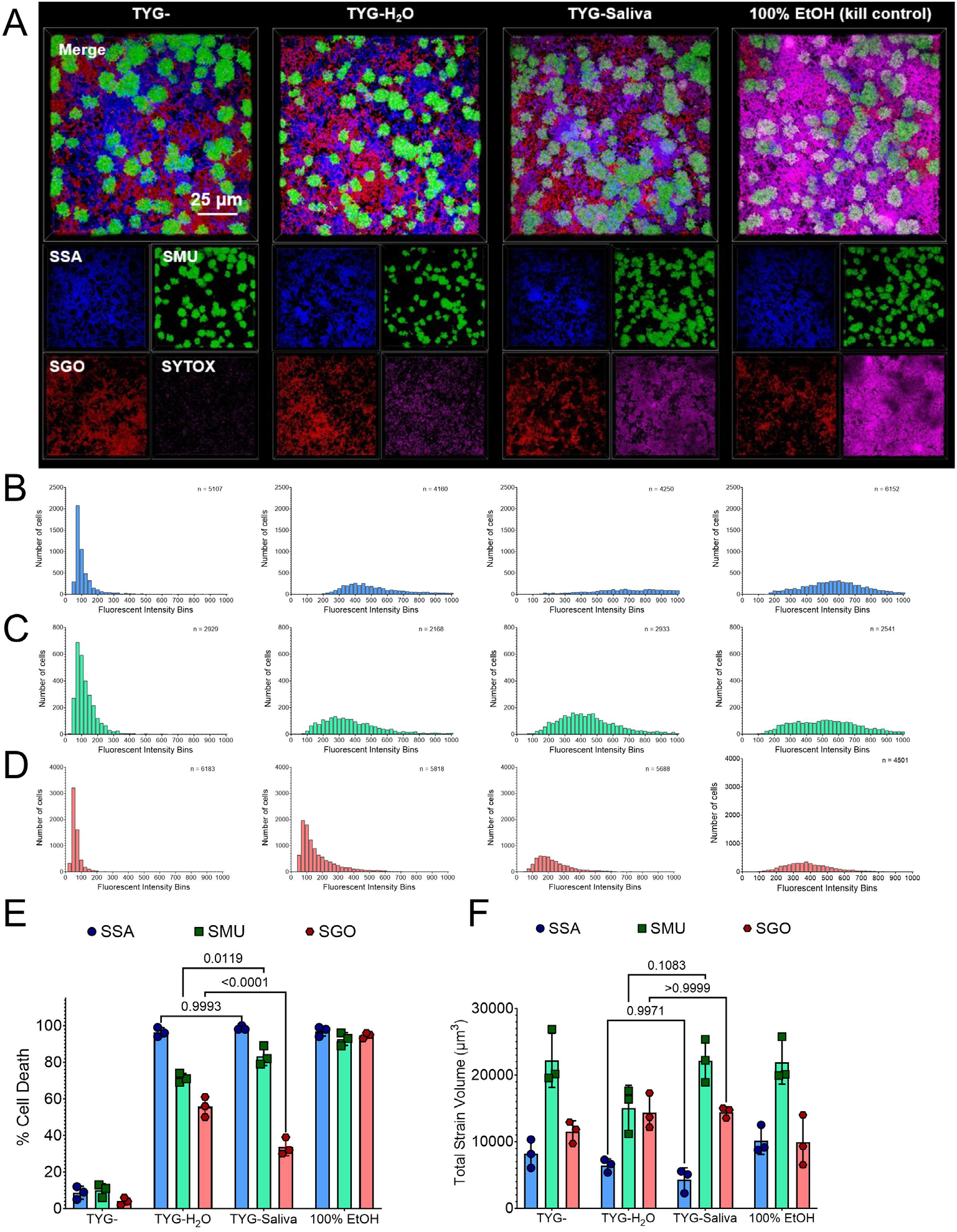
Inclusion of human saliva induces species-specific cell death. **(A)** Maximum intensity, 100x 3D models of a super-resolution confocal-captured biofilm image oriented from the top down (Z+) of *S. sanguinis* (SSA; mTagBFP2, blue), *S. mutans* (SMU; sfGFP, green), and *S. gordonii* (SGO; mScarlet-I3, red) tricultures grown in TYG-, TYG-H_2_O, TYG-Saliva or TYG-with 100% ethanol (EtOH) applied for 15 minutes as a killed control. Biofilms were grown for 24 h with 5 mM sucrose prior to imaging. SYTOX Red dead cell stain (SYTOX, pink) was applied to biofilms prior to imaging. The merged image is shown on top, with individual channels on the bottom. Scale bar (25 µm) is shown in the bottom right of the top merged image. Images are 127 μm (L) x 127 μm (W) x 30 μm (H). Histograms of binned relative fluorescent intensity (a.u.) of the SYTOX dead stain within individual cells from the images shown in A for either **(B)** *S. sanguinis* mTagBFP2, **(C)** *S. mutans* sfGFP or **(D)** *S. gordonii* mScarlet-I3 strains. Each histogram represents the fluorescent intensity from either the (left to right) TYG-, TYG-H_2_O, TYG-Saliva or TYG-with 100% ethanol (EtOH) applied image sets. The height of each bar represents the number of cells (y-axis) within each fluorescent intensity bin. The number of cells analyzed for each graph is shown in the upper right corner. **(E)** Percentage of cell death in the population of either *S. sanguinis* (SSA, blue circles), *S. mutans* (SMU, green squares), or *S. gordonii* (SGO, red hexagons) within each growth condition. The fluorescent intensity threshold to be counted as dead was set at 200 a.u. (determined from the lowest values obtained in the kill control). **(F)** Bar graph of calculated strain volumes of the biofilms shown in A. Three independent images/areas were quantified from the same biofilm sample. Statistical analysis is shown using a two-way analysis of variance (ANOVA) with Tukey’s multiple comparisons test.

To take advantage of the extra channel not occupied by bacterial fluorescence, we also stained the biofilms with SYTOX Red cell stain prior to imaging and quantified the amount of cells that had lost membrane integrity within each species under each condition **(Figure 6B-6E)**. This was achieved by utilizing Nikon’s general analysis program to define cell parameters using artificial intelligence (AI) cell segmentation and by creating a binary mask whereby the fluorescent intensity of each channel could be quantified on a single cell basis. The fluorescent intensity for thousands of cells within each image was first segregated at the species level based on their production of either mTagBFP2, sfGFP or mScarlet-I3, and then binned for the fluorescent intensity of the Alexa647 channel, corresponding to the SYTOX Red stain. We also included a TYG-biofilm that was treated with 100% ethanol (EtOH) for 15 minutes prior to addition of the SYTOX Red stain as a killed control that allowed us to set a threshold value for Alexa647 to count a cell as positive for loss of membrane integrity. 200 a.u. was chosen for this binary threshold value, whereby 92-97% of cells within each species were tallied as positive within the kill control. This was in stark contrast to TYG-without EtOH treatment, where only 4-10% of the cells were positive for SYTOX Red staining. However, in TYG-H_2_O or TYG-Saliva conditions, a large increase in SYTOX staining was observed in a species-specific manner. For example, 96±2% of *S. sanguinis* cells were counted positive in TYG-H_2_O and 99±1% in TYG-Saliva. For *S. gordonii*, those rates were only 56±4% and 34±4%, respectively. *S. mutans* trends more closely followed *S. sanguinis* with 71±2% positive in TYG-H_2_O and 83±4% in TYG-Saliva. The differences in both *S. gordonii* and *S. mutans* between TYG-H_2_O and TYG-Saliva were statistically significant. While there were some observable trends in calculated total strain volumes, such as a decrease in *S. sanguinis* volume from TYG-to TYG-Saliva and a concurrent increase in *S. gordonii* volumes, they were not statistically significant **(Figure 6F)**. In all, this imaging experiment confirms that our constructed strains can be used in mixed-species applications while also examining individual species or strain behaviors at the single-cell level, while displaying unique characteristics of each oral streptococci species during biofilm growth in human saliva that will need to be further pursued.

## DISCUSSION

The study of intermicrobial interactions in the oral cavity has continued to be of interest due to the increased availability of advanced microbiology techniques, including the imaging applications described here. There is still a need to develop tools that increase our capabilities for species, strain, and even single cell-level read-outs. Our goal for this project was to develop a new suite of fluorescent strains that provide a more stable and brighter signal, based on our previous experiences with fluorescent strain development (13). While strains producing GFP have been commonly used in the oral microbiology field with success (28–31), applications with red fluorescent proteins have generally struggled mainly due to signal intensity or brightness (32, 33). Here, we found use of a newly developed mScarlet-I3 (22) extremely successful in producing a strong, channel-specific signal that can be utilized in oral streptococci. In addition, use of blue fluorescent proteins has not been widely adopted within the genus *Streptococcus*. Our successful cloning and application of mTagBFP2 shows that this group of fluorescent proteins can also be used to either “mark” or “tag” cells within biofilms or for other imaging experiments, or potentially used to monitor readouts of promoter activity via promoter fusion constructs. For each species, both mScarlet-I3 and mTagBFP2 displayed fluorescent activities that were recorded higher than sfGFP (Figure 3). There are now over a 1000 different fluorescent proteins listed in FPbase (fpbase.org), many of which are derivatives of the same “ancestor” fluorescent protein that have undergone targeted and/or random mutagenesis to produce a desired result, as the case for mScarlet-I3. We chose to work with both mTagBFP2 and mScarlet-I3 based on their recorded high brightness and their narrow excitation/emission ranges that would make them suitable for imaging multiple fluorescent-producing strains at a time (34). There may still be other blue or red fluorescent proteins that would behave similarly or even better for streptococci imaging, or these proteins could be further developed in the future where new versions could be tested that would improve on the strains described here. We will continue to monitor the fluorescent protein space and develop new strains with desired traits as the need arises or derivatives become available.

While all three stains of *S. mutans* and *S. gordonii* displayed high and strain-specific intensity, we were not able to have similar success with the same constructs in *S. sanguinis*. While we were able to confirm through PCR and sequencing that *S. sanguinis* contained our gene fragment without any known mutations (data not shown), the fluorescent signal derived from these strains was notably weaker for mTagBFP2 and mScarlet-I3, and not detectable over background with sfGFP (Figure 3). While the mTagBFP2 and mScarlet-I3 strains could still be used for imaging applications through user adjustment of instrument settings, we would not suggest further use of sfGFP. These issues may have arose for several reasons, including lack of codon optimization of the coding sequence for *S. sanguinis*. All three fluorescent gene sequences were codon optimized for *S. mutans* prior to gene fragment synthesis, however further optimization may be required in *S. sanguinis* or other species. It was notable that fluorescent intensity for mTagBFP2 and mScarlet-I3 rose after 15 and 18 h, respectively, in *S. sanguinis* compared to 9 h for *S. mutans* and *S. gordonii*, potentially suggesting an issue with protein maturation in this species. We encountered similar issues in our prior work with fluorescent genes encoded on the vector pDL278 (13). It is a challenge to have constructed vectors and/or synthetic gene fragments that do not result in success with every *Streptococcus* species tested, but should be acknowledged that future development of strains may require strategies specific for the species or even the strain an end user has interest in studying.

Issues with *S. sanguinis* were not our only difficulties encountered during this project. We also attempted to clone and test strains with a gene fragment that contained mMaroon1, a far red fluorescent protein with Em/Ex of 609/657 that could be used for visualization in Cy5/Alexa647 imaging channels (35). Similar to the issues with sfGFP in *S. sanguinis*, fluorescent intensity of mMaroon1 could not be quantified over background in any of the three species, nor visualized through either widefield or confocal microscopy (data not shown). A drawback in utilization of far red fluorescent proteins is their lack of brightness, so this result was not entirely surprising. This is another area we will continue to evaluate for further development.

An advancement made with these strains is the ability to use them to monitor single cell-level readouts. This was achievable through signal stability and uniform intensity across the entire population for a given species/strain. Efficient, individual cell segmentation is now practical and allows for monitoring of fluorescent intensity in other channels. As a proof of concept, we observed the amount of cells that displayed loss of cell membrane integrity that allows the SYTOX dye to become penetrant and bind nucleic acid (36, 37), after 24 h of growth for either *S. gordonii, S. mutans*, and *S. sanguinis* in media containing or lacking human saliva. While we did not expect to observe such a drastic change in membrane integrity in media containing human saliva, the experiment shows that strain level data can be collected – both *S. mutans* and *S. sanguinis* saw higher amounts of SYTOX staining than *S. gordonii* grown in the same biofilm. This allows collection of data that would be equivalent to that produced from flow cytometry (i.e., percentages of cells stained or producing a signal within a given population using signal intensity as a read-out), but from an imaging input of cells structured within a biofilm that do not need to be disturbed. This type of data collection can be additionally used in other applications, such as to monitor promoter activities or used in combination with other labeling approaches such as those that utilize click chemistry (38). A limitation for the user may be in data analysis of acquired images; however, we have used built-in Nikon program analysis tools for this study that should be common to similar instruments used in other academic or industry settings.

The availability of these strains, as well as the described blueprint for their design and optimization to construct similar tools in other organisms, will allow us to conduct more specific assays and measurements of mixed-species biofilms for oral streptococci. Use of these tools are not without their limitations – use of fluorescent proteins requires the presence of oxygen in the growth environment and cannot be cultured anaerobically (39), and insertion of the gene fragment is strain specific, requiring further cloning applications to move the gene fragments into other strains of interest. There may be better alternatives, such as fluorescent *in situ* hybridization (FISH) or strain-specific labeling using labeled antibodies/antisera, which are dependent on the user’s experimental goals and applications. However, several advantages such as the ability to make multiple measurements of the same biofilm over time in time course assays as well as the ability to image intact biofilms without the need for fixing, permeabilization or multiple washing steps ensure that these strains will be of use to the field as we continue to explore biofilm formation, community assembly and interspecies interactions between members of the human microbiome.

## MATERIALS AND METHODS

### Strain Inoculation and Growth Medias

Overnight cultures of the bacterial strains used in this study **(Table S1)** were inoculated from single, isolated colonies on Bacto Brain Heart Infusion (Difco BHI; Fisher Bioreagents 237500) agar plates (Difco Agar, Fisher Bioreagents 214010) into BHI broth and incubated at 37°C and 5% CO_2_ with the appropriate antibiotics. Antibiotics were added to overnight growth medium at 1 mg mL^-1^ for both kanamycin and spectinomycin and 0.01 mg mL^-1^ for erythromycin. The next day, cultures were harvested by centrifugation, washed to remove all traces of overnight growth medium, and normalized to OD_600 nm_ = 0.2 with 1X phosphate-buffered saline (PBS) prior to back dilution (1:100) into tryptone and yeast extract supplemented with glucose (TYG, 20 mM glucose final concentration; 10 g tryptone [Fisher Bioreagents BP1421], 5 g yeast extract [Fisher Bioreagents BP1422], 3 g K_2_HPO_4_ [Sigma-Aldrich P3786], and 3.6 g glucose per 1 L H_2_O [Sigma-Aldrich G8270]) media. 1.7 g L^-1^ sucrose (Sigma-Aldrich S7903) was added for all biofilm-related experiments (TYGS; 5 mM sucrose final concentration). Commercially available pooled human saliva was purchased from Innovative Research (IRHUSL250ML). Upon receipt, the saliva was thawed, centrifuged at 4500 RPM for 10 minutes, and then passed through a 0.22 μm filter unit prior to local frozen storage as 10 mL aliquots. For experimental use, the saliva was thawed and used same day. TYG(S) media containing –H_2_O or –Saliva were prepared using protocols detailed elsewhere (27). All strains were maintained for long-term storage at −80ºC in BHI containing 25% glycerol.

### Selection of Gene Fragment Insertion Site and Cloning

Sites for chromosomal insertion of a gene fragment were selected based on a previously published RNA-Seq dataset (18) from mono- and cocultures of strains *S. mutans, S. gordonii* and *S. sanguinis* grown in TYG media until mid-exponential growth phase (OD_600_ ∼ 0.4 – 0.5). The datasets are available from NCBI-GEO (Gene Expression Omnibus) under accession number **GSE209925**. Insertion sites were selected based on read counts at or near 0 within called open reading frames (ORFs) in multiple conditions (mono- and coculture growth), as well as through visual inspection for a lack of reads that mapped within these regions using Integrative Genomics Viewer (IGV, v 2.8.13) (40).

Mutants of the insertion sites were constructed using a PCR ligation mutagenesis approach as previously described (20). Briefly, two gene-flanking proximal sequences corresponding to 750-1000 bp upstream or downstream of the gene of interest were amplified with primer sets containing BamHI restriction sites, digested with BamHI-HF (New England Biolabs, R3136) overnight at 37°C, and ligated to *ermB* (T4 DNA Ligase, New England Biolabs, M0202) (41). The ligation product (∼0.1 µg) was transformed into the strain of interest using a species-specific 0.1 µM final concentration of synthetic CSP peptide (sCSP, Biomatik^™^) within BHI medium and plating the culture onto BHI agar plates containing 0.01 mg/mL erythromycin. Primers used in this study are listed in **Table S2**.

Gene fragments, flanked by BamHI restriction sites, containing a constitutively-active and displaying strong expression in oral streptococci promoter P*veg* (13), either *mtagbfp2* (21), *sfgfp* (15), or *mscarlet-I3* (22) coding sequences that were codon optimized for *S. mutans*, the *aad9* gene conferring spectinomycin resistance (42–44), and the strong transcriptional terminator L3S2P21 (23), were obtained from Integrative DNA Technologies (IDT) and reconstituted in nuclease-free water according to the supplier’s provided protocol. The sequences of each individual component within the gene fragments are listed in **Table S3**, with the entire gene fragment sequence shown in **Supplemental Figures 1-3**. Each gene fragment was amplified with the ‘genefrag-amplify’ primer set prior to undergoing a restriction digest with BamHI-HF, and ligated to two gene-flanking proximal sequences as detailed above. Ligation products where then transformed into either *S. mutans, S. gordonii* or *S. sanguinis* as described above.

### Growth Measurements

Growth measurements were completed using a Bioscreen C MBR automated turbidometric analyzer (Growth Curves Ab Ltd., Helsinki, Finland) with the optical density at 600 nm (OD_600 nm_) recorded every 1 h for 18 h. Wells were overlaid with 0.05 mL sterile mineral oil (Fisher Bioreagents O121). All experiments were completed with three biological replicates measured in technical triplicates.

### Colony Forming Unit Competitive Index Assays

Strains were inoculated according to **Table S4**. Part of the initial inoculum was serially diluted and plated onto two of BHI (selection of WT strains), BHI kanamycin (selection of *S. mutans* pMZ-) or BHI erythromycin (selection of insert site mutants) agar plates. Colony forming units (CFUs) were later enumerated from these agar plates after incubation to determine the initial cell count (t_i_ = 0 h). The remaining inoculum was incubated at 37°C and 5% CO_2_ for 24 h. Resulting cultures were harvested, washed and resuspended with 1X PBS while transferring to a 5 mL polystyrene round-bottom tube. To isolate single cells, tubes were sonicated within a water bath sonicator (four intervals of 30 s, resting 2 min on ice). Cultures were serially diluted and plated onto two of BHI, BHI kanamycin or BHI erythromycin agar plates and incubated for 48 h at 37°C and 5% CO_2_. Following CFU counting of the final cell count (t_f_ = 24 h), a competitive index was calculated using the following formula: CI = Log10 ([t_f_ Wild-type strain or *S. mutans* pMZ-CFU / t_f_ commensal streptococci or insert site mutant CFU] / [t_i_ Wild-type strain or *S. mutans* pMZ-CFU / t_i_ commensal streptococci or insert site mutant CFU]).

### Crystal Violet Biofilm Biomass Quantification

Strains were inoculated into TYGS and incubated for 24 h at 37ºC and 5% CO_2_. Following, medium from the biofilms was aspirated and plates were dunked into a bucket of water to remove all non-attached cells. After drying, 0.05 mL of 0.1% crystal violet (CV; Fisher Chemical C581) was added to each well and incubated at room temperature for 15 minutes. The CV solution was then aspirated and plates were dunked into a bucket of water again to remove excess CV. Plates were dried and imaged. Next, 0.2 mL of 30% acetic acid solution (RICCA Chemical 1383032) was added to extract the bound CV. Extracted CV solution was diluted 1:4 with water into a new 96 well plate before the absorbance at 575 nm was recorded within an Agilent Biotek Synergy H1 multimode plate reader using Gen5 microplate reader software [v 3.11 software]. All biofilm experiments were completed with at least two biological replicates measured in technical quadruplicates.

### Measurements of Strain Fluorescent Intensity

Cultures were plated in TYG along with a 0.05 mL sterile mineral oil overlay and incubated for 24 h at 37ºC in a Agilent Biotek Synergy H1 multimode plate reader with the optical density at 600 nm (OD_600 nm_) and the fluorescent intensity corresponding to DAPI (excitation 399 nm, emission 455 nm, optics bottom, gain 100), FITC (excitation 485 nm, emission 528 nm, optics bottom, gain 100), and Texas Red (excitation 550 nm, emission 590 nm, optics bottom, gain 100) recorded every 0.5 h. For data analysis, the OD_600 nm_ of a medium-only blank was subtracted from respective optical density readings and fluorescent intensities (arbitrary units; a.u.) of wild-type cultures were subtracted from cultures containing fluorescent genes (removing medium/cell background fluorescence). After plotting the resulting data points in GraphPad Prism, an Area Under the Curve (AUC) of the fluorescent intensity was calculated using built-in analysis tools. All experiments were completed with three biological replicates measured in technical quadruplicates.

### Biofilm Microscopy

Bacterial strains were inoculated into TYGS and respective media that contained 1 µM Alexa Fluor 647-labeled dextran (10,000 molecular weight; anionic, fixable; Invitrogen, D22914), added to Cellvis 12 well, glass bottom, black plates (P12-1.5H-N) and incubated at 37°C and 5% CO_2_ for 24 h. Resulting biofilms were first washed with 1X PBS to remove loosely-bound cells and incubated with BSA blocking buffer at room temperature for 0.5 h (Thermo Scientific, 3% BSA is PBS; J61655.AK). Biofilms were then probed with a murine monoclonal antibody against dsDNA (Anti-dsDNA, 3519 DNA, Abcam, ab27156) [2 µg mL^-1^] within BSA blocking buffer for 1 h at room temperature. The biofilms were then washed once and incubated for 1 h at room temperature with an Alexa Fluor 594-labeled goat anti-mouse IgG highly cross-absorbed secondary antibody (Invitrogen, A32742)) [2 µg mL^-1^] within BSA blocking buffer. Finally, the biofilms were washed and stained with Hoechst 33342 solution (5 µM final concentration, Thermo Scientific, 62249) for 15 minutes (if desired).

For widefield microscopy experiments, biofilms were imaged within 1X PBS using a 40X (plan fluorite, 2.7 mm working distance, 0.6 numerical aperture) phase objective on a Agilent Biotek Lionheart FX automated microscope (Agilent Biotek, Winooski, Vermont, United States) equipped with 365 nm, 465 nm, 523 nm and 623 nm light-emitting diodes (LED; 1225007, 1225001, 1225003, 1225005) for acquiring fluorescent signals with DAPI (377/447; 1225100), GFP (469/525; 1225101), RFP (531/593; 1225103) and Cy5 (628/685; 1225105) filter cubes, respectively. Images were captured using Gen5 Image+ software, and quantification of biomass and biofilm thickness were completed either with the Gen5 Image+ software, or by importing .TIF files into BiofilmQ (45). For analysis, single channel images were analyzed by setting object threshold intensity to greater than or equal to 5000 a.u. (arbitrary units) and minimum object size to greater than 5 microns. Options selected included ‘split touching objects’ and ‘fill holes in masks’. Primary edge objects were excluded from analysis. At least four images of each sample, taken at 2500 micron increments to avoid observer bias, were acquired during each experiment.

Confocal imaging was completed with a Nikon A1R system, with resulting image files saved as .nd2 files. Biofilms were visualized by the A1R with the 100X oil objective (1.45 N.A., WD 0.13 mm) using a maximum of 4 channels (DAPI 402 425-475 nm, GFP 487 500-550 nm, Texas Red 561 570-620, and Alexa647 638 663-738 nm). Resonant scanner (x-axis frequency 7.8 kHz, maximum pixel size 512 × 512) was also used. Acquired images were captured as a z-stack with the dimensions of 127.28 × 127.28 μm, a thickness of 30.00 μm, and a step height of 0.50 μm. Imaging processing was completed with Nikon NIS-Elements software (v 5.22.00). Image quality and clarity was improved using the NIS-Elements Denoise.ai feature across all samples and prior to further analysis such as quantified volume calculations for each strain/component. Maximum intensity and z-depth coded 3D images were derived from these normalized, denoised images within NIS-Elements.

### SYTOX Red Staining and Single Cell Analysis

Prior to imaging, biofilms were stained with the SYTOX Red cell stain (Invitrogen, S34859) for 15 minutes at room temperature, according to the supplier’s protocol. The stain was then removed, biofilms washed and imaged within 1X PBS.

For single cell analysis, NIS-Elements General Analysis (GA3), in combination with the existing NIS-Elements software AI tool, was utilized. Custom thresholds for each channel were developed and applied uniformly across all images. This procedure consisted of the creation of a binary mask to define cell parameters, along with AI cell segmentation so further analysis could be done on the volume and shape of the defined cell structures. Alexa647 channel intensities, corresponding to the SYTOX Red cell stain, could then be measured within segmented binary cell structures. Acquired data was exported to excel and later analyzed in GraphPad Prism.

### Graphing and Statistics

Graphing of data was completed with GraphPad Prism [version 10.1.2] software. All statistical analysis was completed within GraphPad Prism using the built-in analysis tools, including unpaired t-tests with Welch’s correction, one-way or two-way ANOVAs with post hoc tests (Sidak’s, Dunnett’s or Tukey’s test, respectively) for a multiple comparison, and AUC calculations.

## ACKNOLWEDGEMENTS

This work was supported by a grant from the National Institute of Dental and Craniofacial Research (NIDCR) of the National Institutes of Health (NIH) R03DE031766. Super-resolution confocal imaging was completed with resources from the Campus Microscopy and Imaging Facility (CMIF) and the OSU Comprehensive Cancer Center (OSUCCC) Microscopy Shared Resource (MSR), The Ohio State University. This facility is supported in part by grant P30 CA016058, National Cancer Institute, Bethesda, MD.

## AUTHOR CONTRIBUTIONS

<u>Daniel Peters</u> – cloned strains, collected and performed data analysis, contributed to methodology development, and participated in review and editing of the manuscript.

<u>Iris Shin</u> – cloned strains, collected and performed data analysis and participated in review and editing of the manuscript.

<u>Alyssa Deever</u> -- collected and performed data analysis and participated in review and editing of the manuscript.

<u>Justin Kaspar</u> - conceptualized project, collected and performed data analysis, contributed to methodology development, wrote the original draft and participated in review and editing of the manuscript.

## DECLARATION OF CONFLICTING INTERESTS

The authors declared no potential conflicts of interest with respect to the research, authorship, and/or publication of this article.

## Notes

### Competing Interest Statement

The authors have declared no competing interest.

