## Supplemental Figures 1-3 for "Design, Development and Validation of New Fluorescent Strains for Studying Oral Streptococci"

### Table of Contents:

Supplemental Figures (S1-S4): Pages 2 - 5

Supplemental Tables (Table S1-S4): Pages 6 – 11

Supplemental References: Page 12

[#] Corresponding author

**Mailing address:**
Division of Biosciences, The Ohio State University, College of Dentistry,
305 W. 12th Avenue, Postle Hall Rm 4185, Columbus, OH 43210.


cagcaaagaatggcggaaacgtaaaagaagttatggaaataagacttagaagcaaacttaagagtgtgtt
gacagtgcagtaccttaaaatttgtataataggaattgaagttaaattagatgctaaaatttt**GGATCC**
aagaaggagtgattacGAGCTCTAGATCGAATTCCTTATTAACGTTGATATAATTTAAATTTTATTTGAC
AAAAATGGGCTCGTGTTGTACAATAAATGTGATTAACTAATAAGGAGGACAAACatgagcgagctgatta
aggagaacatgcacatgaagctgtacatggagggcaccgtggacaaccatcacttcaagtgcacatccga
gggcgaaggcaagccctacgagggcacccagaccatgagaatcaaggtggtcgagggcggccctctcccc
ttcgccttcgacatcctggctactagcttcctctacggcagcaagaccttcatcaaccacacccagggca
tccccgacttcttcaagcagtccttccctgagggcttcacatgggagagagtcaccacatacgaagacgg
gggcgtgctgaccgctacccaggacaccagcctccaggacggctgcctcatctacaacgtcaagatcaga
ggggtgaacttcacatccaacggccctgtgatgcagaagaaaacactcggctgggaggccttcaccgaga
cgctgtaccccgctgacggcggcctggaaggcagaaacgacatggccctgaagctcgtgggcgggagcca
tctgatcgcaaacgccaagaccacatatagatccaagaaacccgctaagaacctcaagatgcctggcgtc
tactatgtggactacagactggaaagaatcaaggaggccaacaacgagacctacgtcgagcagcacgagg
tggcagtggccagatactgcgacctccctagcaaactggggcacaagcttaattaaaacgtaaaagaagt
taatgaggaggatatatttgaatacatacgaacaaattaataaagtgaaaaaaatacttcggaaacattt
aaaaaataaccttattggtacttacatgtttggatcaggagttgagagtggactaaaaccaaatagtgat
cttgacttttttagtcgtcgtatctgaaccattgacagatcaaagtaaagaaatacttatacaaaaaatta
gacctatttcaaaaaaaataggagataaaagcaacttacgatatattgaattaacaattattattcagca
agaaatggtaccgtggaatcatcctcccaaacaagaatttatttatggagaatggttacaagagctttat
gaacaaggatacattcctcagaaggaattaaattcagatttaaccataatgctttaccaagcaaaacgaa
aaaataaaagaatatacggaaattatgacttagaggaattactacctgatattccattttctgatgtgag
aagagccattatggattcgtcagaggaattaatagataattatcaggatgatgaaaccaactctatatta
actttatgccgtatgattttaactatggacacgggtaaaatcataccaaaagatattgcgggaaatgcag
tggctgaatcttctccattagaacatagggagagaattttgttagcagttcgtagttatcttggagagaa
tattgaatggactaatgaaaatgtaaatttaactataaactatttaaataacagattaaaaaattataa
taaCTCGGTACCAAATTCCAGAAAAGAGGCCTCCCGAAAGGGGGGCCTTTTTTCGTTTTGGTCCttctat
**GGATCC**ctttttgtaaatttggaaagttacacgttactaaagggactgtagatccagcaggtatactactg
acagc

**Supplemental Figure 1. Overview of P*veg*:*mtagbfp2* gene fragment design.** Sequence of the P*veg*:*mtagbfp2* gene fragment that was synthesized and used in cloning for all mTagBFP2 strains. Several features are either highlighted or in colored text: Location of the genefrag-amplify primers is shown in yellow highlight, the BamHI restriction sites are shown in **turquoise highlight and bolded**, the Pveg promoter is shown in PINK CAPITAL letters, the *mtagbfp2* gene is shown in blue letters, *aad9* sequence is shown in light gray letters, and the transcriptional terminator, L3S2P21, is shown in PURPLE CAPITAL letters. The gene fragment was synthesized by Integrated DNA Technologies (IDT).

cagcaaagaatggcggaaacgtaaaagaagttatggaaataagacttagaagcaaacttaagagtgtgtt
gacagtgcagtaccttaaaattttgtataataggaattgaagttaaattagatgctaaaatttt**GGATCC**
aagaaggagtgattacGAGCTCTAGATCGAATTCCTTATTAACGTTGATATAATTTAAATTTTATTTGAC
AAAAATGGGCTCGTGTTGTACAATAAATGTGATTAACTAATAAGGAGGACAAACatgtcaaaggagaag
agctgttcacaggtgttgtgccgattctcgttgagcttgacggagatgtaaacggacacaaattctctgt
tcgcggtgaaggtgaaggagatgcaacaaacggcaagctgacattgaagtttatttgcacaactggaaag
ctgccggttccttggccgacacttgtaacgacgctgacttacggcgttcaatgcttctctcgttatccag
accacatgaaacgccatgatttcttcaaatctgcaatgcctgaaggctacgttcaagagcgtacgatcag
cttcaaagatgacggaacgtacaaaacaagagcagaagtgaagtttgaaggtgacacacttgtgaaccgc
attgaattgaaaggcattgatttcaaagaagatggaaacatccttggacacaaacttgaatacaacttca
acagccacaacgtatacatcactgctgacaaacaaaaaaacggcatcaaagcaaacttcaaaatccgtca
taacgtagaggacggttctgttcagcttgctgatcattatcagcaaaatacaccgatcggtgacggcccg
gttcttcttcctgataaccattatttatcaactcaaagcgtattatcaaaagacccaaatgaaaagcgtg
accacatggtgctgcttgaatttgtgacagctgctggtatcactcacggcatggatgagctttataagta
atttggaaagttacacgatgagacgcatttaccttaatacatatgagcagatcaacaaggtgaagaagat
tttaagaaagcacttaaaaaataatcttattggcacgtatatgttcggaagcggtgtcgaatcaggtctt
aagccgaattctgacttagacttcttggtcgttgtctctgagcctttaacggaccaatctaaagaaattt
tgattcaaaaaattcgccctatctcaaagaaaatcggtgacaagtcaaatttgagatacattgaattaac
catcatcatccagcaagaaatggtcccgtggaaccacccgccgaagcaagagttcatttacggcgaatgg
ttacaggagttgtatgagcaaggctacattccacagaaagagcttaatagtgacttgacaatcatgttat
atcaggcaaaacgtaagaacaaacgcatttacggaaactatgatttagaggaacttttgcccgatatccc
atttctgacgttcgtcgcgccattatggacagctctgaggagttaattgataactaccaggatgatgag
acgaatagtattttaactctttgtcgtatgattttgacaatggacactggtaaaatcatccccaaggata
ttgctggtaatgccgttgcagaaagcagcccattggagcacagagagcgcattcttcttgcagtacgcag
ctatcttggagagaatatcgagtggacaaacgagaatgttaatttgacaatcaattatttgaacaatcgc
ttgaaaaagctttaataaCTCGGTACCAAATTCCAGAAAAGAGGCCTCCCGAAAGGGGGGCCTTTTTTCG
TTTTGGTCCttctat**GGATCC**cttttgtaaatttggaaagttacacgttactaaagggactgtagatcca
gcaggtatactactgacagc

**Supplemental Figure 2. Overview of P*veg*:*sfgfp* gene fragment design.** Sequence of the P*veg*:*sfgfp* gene fragment that was synthesized and used in cloning for all sfGFP strains. Several features are either highlighted or in colored text: location of the 'genefrag-amplify' primers is shown in yellow highlight, the BamHI restriction sites are shown in **turquoise highlight and bolded**, the Pveg promoter is shown in PINK CAPITAL letters, the *sfgfp* gene is shown in green letters, *aad9* sequence** is shown in light gray letters, and the transcriptional terminator, L3S2P21, is shown in PURPLE CAPITAL letters. The gene fragment was synthesized by Integrated DNA Technologies (IDT).

**Note: the *aad9* sequence in this gene fragment underwent codon optimization for *S. mutans* using IDT's codon optimizer tool. This was done in order to meet the gene fragment complexity benchmarks during IDT gene block order submission (synthesis of a gene block using the *aad9* sequence included in the other two fragments failed and was re-attempted and successful with this sequence).

```
cagcaaagaatggcggaaacgtaaaagaagttatggaaataagacttagaagcaaacttaagagtgtgtt
gacagtgcagtaccttaaaattttgtataataggaattgaagttaaattagatgctaaaaatttGGATCC
aagaaggagtgattacGAGCTCTAGATCGAATTCCTTATTAACGTTGATATAATTTAAATTTTATTTGAC
AAAAATGGGCTCGTGTTGTACAATAAATGTGATTAACTAATAAGGAGGACAAACatggattctaccgaag
ctgttatcaaagagttcatgcgttttaaggtacacatggagggctcaatgaatggacacgaatttgaaat
tgaaggagagggtgagggtcgcccgtacgaaggcacgcaaacggctaaattgaaagtaacgaaaggcggc
ccgttgccatttagttgggacatcttgtcaccgcaatttatgtatggttcacgcgcttttatcaagcacc
cggccgatattcctgactactggaagcaatcattcccagagggcttcaagtgggagcgcgttatgatctt
cgaggacggtggcacagtctcagttacgcaagacacctctcttgaggacggaactttgatttacaaagtg
aagttgcgtggaggtaatttcccgcccgatggaccggtcatgcagaaacgtaccatgggctgggaggctt
caacagagcgcctttacccggaagatgtcgtacttaaaggcgacatcaagatggcattgcgtttgaaaga
tggtggacgttaccttgccgatttcaagaccacctataaggcaaaaaagccagtgcaaatgcccggcgcc
tttaatattgaccgcaagttggacatcacgagtcacaatgaggactatactgttgttgagcaatatgagc
gttctgttgcaagacattcaacaggtggtagcggcggtagttaagagtgtgttgacagtgatgaggagga
tatatttgaatacatacgaacaaattaataaagtgaaaaaaatacttcggaaacatttaaaaaataacct
tattggtacttacatgtttggatcaggagttgagagtggactaaaaccaaatagtgatcttgacttttta
gtcgtcgtatctgaaccattgacagatcaaagtaaagaaatacttatacaaaaaattagacctatttcaa
aaaaaataggagataaaagcaacttacgatatattgaattaacaattattattcagcaagaaatggtacc
gtggaatcatcctcccaaacaagaatttatttatggagaatggttacaagagctttatgaacaaggatac
attcctcagaaggaattaaattcagatttaaccataatgctttaccaagcaaaacgaaaaaataaaagaa
tatacggaaattatgacttagaggaattactacctgatattccatttctgatgtgagaagagccattat
ggattcgtcagaggaattaatagataattatcaggatgatgaaaccaactctatattaactttatgccgt
atgattttaactatggacacgggtaaaatcataccaaaagatattgcgggaaatgcagtggctgaatctt
ctccattagaacatagggagagaattttgttagcagttcgtagttatcttggagagaatattgaatggac
taatgaaaatgtaaatttaactataaactatttaaataacagattaaaaaattataataaCTCGGTACC
AAATTCCAGAAAAGAGGCCTCCCGAAAGGGGGGCCTTTTTTCGTTTTGGTCCttctatGGATCCcttttg
taaatttggaaagttacacgttactaaagggactgtagatccagcaggtatactactgacagc
```

**Supplemental Figure 3. Overview of P*veg*:*mscarlet-I3* gene fragment design.** Sequence of the P*veg*:*mscarlet-I3* gene fragment that was synthesized and used in cloning for all mScarlet-I3 strains. Several features are either highlighted or in colored text: location of the 'genefrag-amplify' primers is shown in yellow highlight, the BamHI restriction sites are shown in **turquoise highlight and bolded**, the Pveg promoter is shown in PINK CAPITAL letters, the *mscarlet-I3* gene is shown in red letters, *aad9* sequence is shown in light gray letters, and the transcriptional terminator, L3S2P21, is shown in PURPLE CAPITAL letters. The gene fragment was synthesized by Integrated DNA Technologies (IDT).

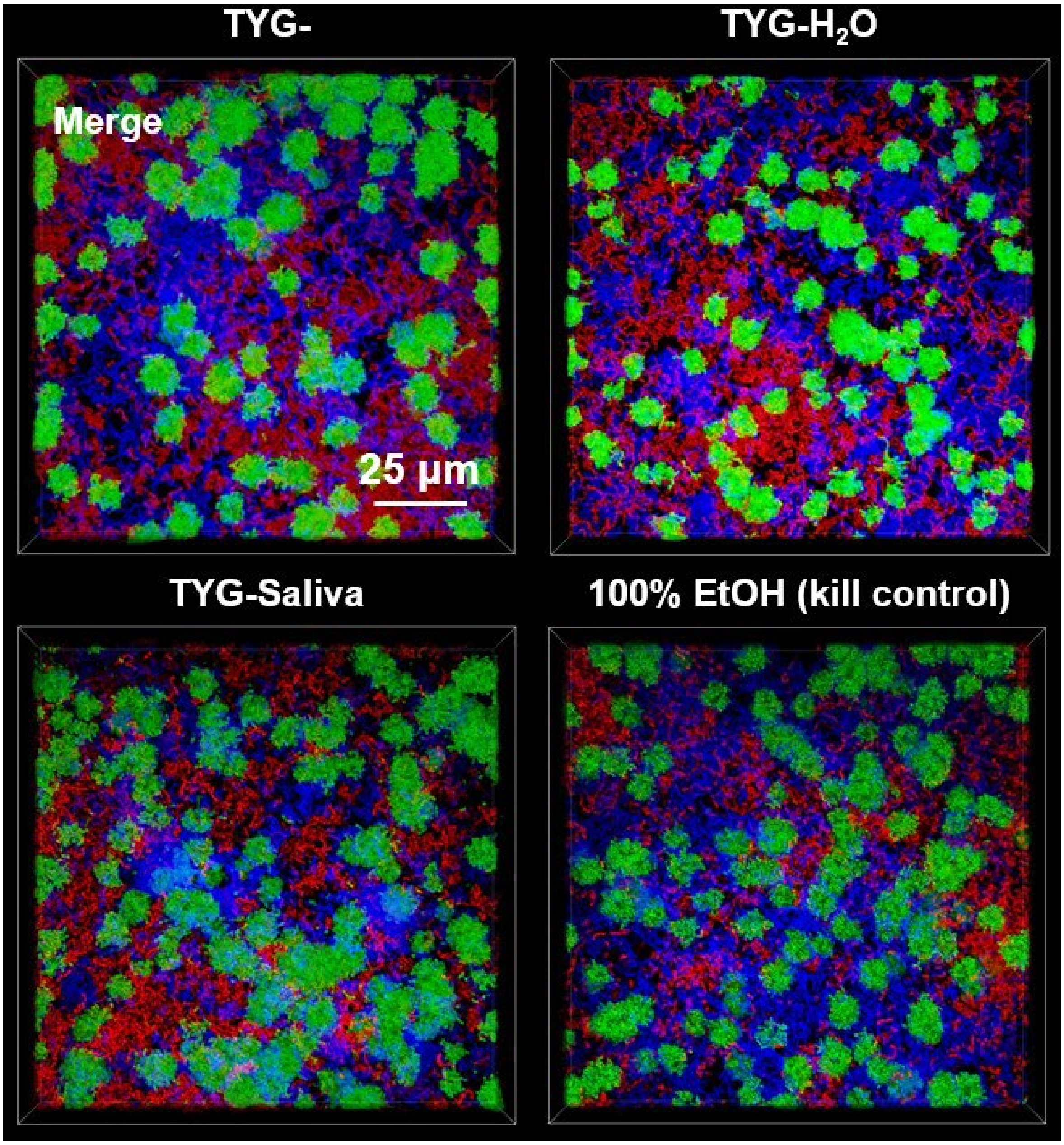


**Supplemental Figure 4. Merged biofilm images from Figure 6 without the SYTOX Red channel overlay.** Maximum intensity, 100x 3D models of a super-resolution confocal-captured biofilm image oriented from the top down (Z+) of *S. sanguinis* (mTagBFP2, blue), *S. mutans* (sfGFP, green), and *S. gordonii* (mScarlet-I3, red) tricultures grown in TYG-, TYG-$H_2O$, TYG-Saliva or TYG- with 100% ethanol (EtOH) applied for 15 minutes as a kill control. Biofilms were grown for 24 h with 5 mM sucrose prior to imaging. These are the same merged images from Figure 6, but without the SYTOX Red (CY5 filter) overlay. Scale bar (25 µm) is shown in the top left merged image. Images are 127 µm (L) x 127 µm (W) x 30 µm (H).

### SUPPLEMENTAL TABLES

**Supplemental Table 1.** Bacterial strains used in this study.

| Species | Strain | Genotype or description | Antibiotic resistance | Reference or source |
|---|---|---|---|---|
| *Streptococcus gordonii* | DL1 | | | Lab Stock |
| | Δ2075 | Allelic exchange of SGO_2075, a gene not transcriptionally active in TYG, with *ermB* providing resistance to erythromycin | Erythromycin | This manuscript |
| | Δ2075::P*veg*:*mtagbfp*2 | Allelic exchange of SGO_2075 with a gene block containing the fluorescent gene *mtagbfp2*, driven by the P*veg* promoter and carrying *aad9* for spectinomycin resistance | Spectinomycin | This manuscript |
| | Δ2075::P*veg*:*sfgfp* | Allelic exchange of SGO_2075 with a gene block containing the fluorescent gene *sfgfp*, driven by the P*veg* promoter and carrying *aad9* for spectinomycin resistance | Spectinomycin | This manuscript |
| | Δ2075::P*veg*:*mscarlet-I3* | Allelic exchange of SGO_2075 with a gene block containing the fluorescent gene *mscarlet-I3*, driven by the P*veg* promoter and carrying *aad9* for spectinomycin resistance | Spectinomycin | This manuscript |
| *Streptococcus mutans* | UA159 | | | Lab Stock |
| | pMZ- / UA159 | UA159 with pMZ plasmid integrated into genome. Serves as a "marked" *S. mutans* strain during colony forming unit (CFU) assays | Kanamycin | (Shields et al. 2019) |
| | Δ1155 | Allelic exchange of SMU_1155, a gene not transcriptionally active in TYG, with *ermB* providing resistance to erythromycin | Erythromycin | This manuscript |

| | | | | |
|---|---|---|---|---|
| | Δ1155::P*veg*:*mtagbfp*2 | Allelic exchange of SMU_1155 with a gene block containing the fluorescent gene *mtagbfp2*, driven by the P*veg* promoter and carrying *aad9* for spectinomycin resistance | Spectinomycin | This manuscript |
| | Δ1155::P*veg*:*sfgfp* | Allelic exchange of SMU_1155 with a gene block containing the fluorescent gene *sfgfp*, driven by the P*veg* promoter and carrying *aad9* for spectinomycin resistance | Spectinomycin | This manuscript |
| | Δ1155::P*veg*:*mscarlet-I3* | Allelic exchange of SMU_1155 with a gene block containing the fluorescent gene *mscarlet-I3*, driven by the P*veg* promoter and carrying *aad9* for spectinomycin resistance | Spectinomycin | This manuscript |
| *Streptococcus sanguinis* | SK36 | | | Lab Stock |
| | Δ2030 | Allelic exchange of SSA_2030, a gene not transcriptionally active in TYG, with ermB providing resistance to erythromycin | Erythromycin | This manuscript |
| | Δ2030::P*veg*:*mtagbfp*2 | Allelic exchange of SSA_2030 with a gene block containing the fluorescent gene *mtagBFP2*, driven by the P*veg* promoter and carrying *aad9* for spectinomycin resistance | Spectinomycin | This manuscript |
| | Δ2030::Pveg:*sfgfp* | Allelic exchange of SSA_2030 with a gene block containing the fluorescent gene *sfgfp*, driven by the P*veg* promoter and carrying *aad9* for spectinomycin resistance | Spectinomycin | This manuscript |
| | Δ2030::Pveg:*mscarlet-I3* | Allelic exchange of SSA_2030 with a gene block containing the fluorescent gene *mscarlet-I3*, driven by the P*veg* promoter and carrying *aad9* for spectinomycin resistance | Spectinomycin | This manuscript |

**Supplemental Table 2.** Primers used in this study.

| Species and Strain | Primer Name | Primer Sequence (5' - 3')* | Tm |
|---|---|---|---|
| | genefrag-amplify-F | GAG TGT GTT GAC AGT GCA GTA CC | 58 |
| | genefrag-amplify-R | CCT GCT GGA TCT ACA GTC C | 55 |
| | genefrag-check-F | GTT TGG ATC AGG AGT TGA GAG TGG | 57 |
| | genefrag-check-R | CTA ATG GAG AAG ATT CAG CCA CTG C | 58 |
| *Streptococcus gordonii* DL1 | SGO_2075_A | TTC CTA TAG GAA TAG GAA CAG GGC | 55 |
| | SGO_2075_B | CTA **<u>GGA TCC</u>** TTT CGG GTG CAC TTT CCT G | 62 |
| | SGO_2075_C | TCT **<u>GGA TCC</u>** ACG TCG GTA TAA CCC | 59 |
| | SGO_2075_D | CTT AGC CGG GTT AGT AGT CAA GCC | 59 |
| *Streptococcus mutans* UA159 | SMU_1155_A | CTC ATA GGA ATC AAC TGG ACA GGC C | 59 |
| | SMU_1155_B | GAT **<u>GGA TCC</u>** CGG GAT CGT CTG GCC | 65 |
| | SMU_1155_C | AAG **<u>GGA TCC</u>** CGC AGC TCT TCC ACC | 65 |
| | SMU_1155_D | TTC CAC AGT GAC CTC AGA AGC TGC | 61 |
| *Streptococcus sanguinis* SK36 | SSA_2030_A | GTC TTC TTG GAC TTG CAG AGC TGC | 60 |
| | SSA_2030_B | GGT **<u>GGA TCC</u>** TGC TTG TTC ATA GCT TGG CG | 64 |
| | SSA_2030_C | CAT **<u>GGA TCC</u>** ATT ATC GGC TAT CCC AG | 58 |
| | SSA_2030_D | TAA GTA ACG TTC CAG AGC CTC CTC | 57 |

***<u>Bold and underline</u>** denotes BamHI cut site used in the PCR ligation mutagenesis approach

The 'genefrag-amplify' primers were used to amplify the entire fluorescent gene fragment synthesized by IDT prior to BamHI restriction digest.

The 'genefrag-check' primers were used during colony PCR to screen for desired transformants after transformation of the linear ligation product. These primers amplify a region within the *aad9* gene.

**Supplemental Table 3.** Sequences and sources of individual components of the fluorescent gene fragments.

| Component | Sequence | Source |
|---|---|---|
| P*veg* promoter | gagctctagatcgaattccttattaacgttgatataatttaaattttatttgacaaaaatgggctcgtg<br>ttgtacaataaatgtgattaactaataaggaggacaaac | (Shields et al. 2019) |
| *aad9* gene | atgaggaggatatatttgaatacatacgaacaaattaataaagtgaaaaaaatacttcggaaa<br>catttaaaaaataaccttattggtacttacatgtttggatcaggagttgagagtggactaaaacca<br>aatagtgatcttgactttttagtcgtcgtatctgaaccattgacagatcaaagtaaagaaatacttat<br>acaaaaaattagacctatttcaaaaaaaaataggagataaaagcaacttacgatatattgaatta<br>acaattattattcagcaagaaatggtaccgtggaatcatcctcccaaacaagaatttatttatgga<br>gaatggttacaagagctttatgaacaaggatacattcctcagaaggaattaaattcagatttaac<br>cataatgctttaccaagcaaaacgaaaaaataaaagaatatacggaaattatgacttagagg<br>aattactacctgatattccattttctgatgtgagaagagccattatggattcgtcagaggaattaata<br>gataattatcaggatgatgaaaccaactctatattaactttatgccgtatgattttaactatggacac<br>gggtaaaatcataccaaaagatattgcgggaaatgcagtggctgaatcttctccattagaacat<br>agggagagaattttgttagcagttcgtagttatcttggagagaatattgaatggactaatgaaaat<br>gtaaatttaactataaactatttaaataacagattaaaaaaattataataa | (Benson and Haldenwang 1993; Guérout-Fleury et al. 1995; LeDeaux and Grossman 1995) |
| L3S2P21 transcriptional terminator | ctcggtaccaaattccagaaaagaggcctcccgaaaggggggcctttttcgttttggtcc | (Chen et al. 2013) |
| *mtagbfp2* gene | atgagcgagctgattaaggagaacatgcacatgaagctgtacatggagggcaccgtggaca<br>accatcacttcaagtgcacatccgagggcgaaggcaagccctacgagggcacccagaccat<br>gagaatcaaggtggtcgagggcggccctctcccсttcgccttcgacatcctggctactagcttcc<br>tctacggcagcaagaccttcatcaaccacacccagggcatccccgacttcttcaagcagtcctt<br>ccctgagggcttcacatgggagagagtcaccacatacgaagacgggggcgtgctgaccgct<br>acccaggacaccagcctccaggacggctgcctcatctacaacgtcaagatcagaggggtga<br>acttcacatccaacggccctgtgatgcagaagaaaacactcggctgggaggccttcaccgag<br>acgctgtaccccgctgacggcggcctggaaggcagaaacgacatggccctgaagctcgtgg<br>gcgggagccatctgatcgcaaacgccaagaccacatatagatccaagaaacccgctaaga<br>acctcaagatgcctggcgtctactatgtggactacagactggaaagaatcaaggaggccaac<br>aacgagacctacgtcgagcagcacgaggtggcagtggccagatactgcgacctccctagca<br>aactggggcacaagcttaattaa | (Subach et al. 2011) |
| *sfgfp* gene | atgtcaaaaggagaagagctgttcacaggtgttgtgccgattctcgttgagcttgacggagatgt<br>aaacggacacaaattctctgttcgcggtgaaggtgaaggagatgcaacaaacggcaagctg<br>acattgaagtttatttgcacaactggaaagctgccggttccttggccgacacttgtaacgacgctg<br>acttacggcgttcaatgcttctctcgttatccagaccacatgaaacgccatgatttcttcaaatctgc<br>aatgcctgaaggctacgttcaagagcgtacgatcagcttcaaagatgacggaacgtacaaaa<br>caagagcagaagtgaagtttgaaggtgacacacttgtgaaccgcattgaattgaaaggcattg<br>atttcaaagaagatggaaacatccttggacacaaacttgaatacaacttcaacagccacaacg<br>tatacatcactgctgacaaacaaaaaaacggcatcaaagcaaacttcaaaatccgtcataac<br>gtagaggacggttctgttcagcttgctgatcattatcagcaaaatacaccgatcggtgacggccc<br>ggttcttcttcctgataaccattatttatcaactcaaagcgtattatcaaaagacccaaatgaaaa<br>gcgtgaccacatggtgctgcttgaatttgtgacagctgctggtatcactcacggcatggatgagct<br>ttataagtaa | (Overkamp et al. 2013) |

| | | |
|---|---|---|
| *mscarlet-I3* gene | atggattctaccgaagctgttatcaaagagttcatgcgttttaaggtacacatggagggctcaatg<br>aatggacacgaatttgaaattgaaggagagggtgagggtcgcccgtacgaaggcacgcaaa<br>cggctaaattgaaagtaacgaaaggcggcccgttgccatttagttgggacatcttgtcaccgca<br>atttatgtatggttcacgcgcttttatcaagcacccggccgatattcctgactactggaagcaatca<br>ttcccagagggcttcaagtgggagcgcgttatgatcttcgaggacggtggcacagtctcagttac<br>gcaagacacctctcttgaggacggaactttgatttacaaagtgaagttgcgtggaggtaatttcc<br>cgcccgatggaccggtcatgcagaaacgtaccatgggctgggaggcttcaacagagcgcctt<br>tacccggaagatgtcgtacttaaaggcgacatcaagatggcattgcgtttgaaagatggtggac<br>gttaccttgccgatttcaagaccacctataaggcaaaaaagccagtgcaaatgcccggcgcct<br>ttaatattgaccgcaagttggacatcacgagtcacaatgaggactatactgttgttgagcaatatg<br>agcgttctgttgcaagacattcaacaggtggtagcggcggtagttaa | (Gadella et al. 2023) |

**Supplemental Table 4.** Concentration of inoculated bacterial strains for competitive index and biofilm triculture experiments.

from (Choi et al. 2024)

| Strain Name | Concentration of Competitor | ***S. mutans*** **Concentration** |
|---|---|---|
| *S. gordonii* DL1 | 10% solution (1:10 dilution); 1x | 1x |
| *S. sanguinis* SK36 | 1x | 1x |
